# Exofacial membrane composition and lipid metabolism regulates plasma membrane P4-ATPase substrate specificity

**DOI:** 10.1101/2020.06.24.169672

**Authors:** Bartholomew P. Roland, Bhawik K. Jain, Todd R. Graham

**Author notes:** These authors contributed equally to the work (Bartholomew P. Roland, Bhawik K. Jain). To whom correspondence should be addressed: Todd R. Graham: Department of Biological Sciences, Vanderbilt University Nashville, TN 37235.

## Abstract

The plasma membrane of a cell is characterized by an asymmetric distribution of lipid species across the exofacial and cytofacial aspects of the bilayer. The regulation of membrane asymmetry is a fundamental characteristic of membrane biology, and is crucial for signal transduction, vesicle transport, and cell division. The type-IV family of P-ATPases, or P4-ATPases, establish membrane asymmetry by selection and transfer of a subset of membrane lipids from the lumenal or exofacial leaflet to the cytofacial aspect of the bilayer. It is still unclear how these enzymes sort through the spectrum of lipids within the membrane to identify their desired substrate(s) and how the membrane environment modulates this activity. Therefore, we tested how the yeast plasma membrane P4-ATPase, Dnf2, responds to changes in membrane composition induced by perturbation of endogenous lipid biosynthetic pathways or exogenous application of lipid. The primary substrates of Dnf2 are two chemically divergent lipids, glucosylceramide (GlcCer) and phosphatidylcholine ((PC) or their lyso-lipid derivatives), and we find that these substrates compete with each other for transport. Acutely inhibiting sphingolipid synthesis using myriocin attenuates transport of exogenously applied GlcCer without perturbing PC transport. Deletion of genes controlling later steps of glycosphingolipid production also perturb GlcCer transport to a greater extent than PC transport. Surprisingly, application of lipids that are poor transport substrates differentially affect PC and GlcCer transport by Dnf2, thus altering substrate preference. Our data indicate that Dnf2 exhibits exquisite sensitivity to the membrane composition; thus, providing feedback onto the function of the P4-ATPases.

## Introduction

The plasma membrane is organized as a lipid bilayer consisting of exofacial and cytofacial leaflets. Different lipids are distributed heterogeneously between these two leaflets to form an asymmetric membrane structure. The exofacial side of the membrane is enriched with phosphatidylcholine (PC) and sphingolipids (sphingomyelin and glycosphingolipids), and the cytosolic side is enriched for phosphatidylserine (PS) and phosphatidylethanolamine (PE) (1–3). This asymmetric nature of lipid bilayer is implicated in different cellular processes, such as vesicular trafficking, apoptosis, signal transduction and cytokinesis. Alteration in lipid asymmetry also plays an important role in physiological functions like blood coagulation and host-pathogen interactions (4).

Membrane asymmetry is generated primarily by ATP-dependent lipid transporters called P4-ATPases, or flippases. These integral membrane enzymes transport lipid substrates from the exofacial to the cytosolic leaflets of the membrane. P4-ATPases are heterodimers with a catalytic α subunit (P4-ATPase) and a noncatalytic β subunit (Cdc50, Lem3 or Crf1 in yeast, CDC50A, CDC50B or CDC50C in humans). The budding yeast *Saccharomyces cerevisiae* expresses 5 P4-ATPases: Dnf1, Dnf2, Dnf3, Drs2 and Neo1; while 14 different P4-ATPases are expressed in humans. (5–7). Mutations in these human P4-ATPases are associated with neurological disease, cholestasis, reproductive dysfunction, and metabolic disease (8, 9)

Understanding the substrate specificity of these transporters is essential for determining their role in health and disease. P4-ATPases were first described to be aminophospholipid (PS, PE) translocases and have traditionally been thought to specifically transport glycerophospholipids from the exofacial to the cytofacial side of the bilayer (10). Recently, however, we and others discovered that a group of yeast and human P4-ATPases translocate the sphingolipid glucosylceramide (GlcCer). GlcCer is the primary substrate for the human P4-ATPases ATP10B and ATP10D, while ATP10A transports both PC and GlcCer (11, 12). GlcCer is a central intermediate in sphingolipid biosynthesis and acts as a precursor for the synthesis of all complex glycosphingolipids, such as gangliosides and globosides in animal cells (13). In humans, GlcCer accumulation has been associated with Gaucher and Parkinson’s diseases. Mutations in the endo/lysosomal ATP10B are also linked to Parkinson’s disease, cause accumulation of GlcCer, lysosomal dysfunction and loss of cortical neurons (12). In addition, mutations in ATP10A and ATP10D are associated with diet-induced obesity, insulin resistance, myocardial infarction and atherosclerosis (14–19). Thus, GlcCer metabolism and transport play crucial roles in various pathologies.

The GlcCer flippases in yeast are Dnf1 and Dnf2, orthologs of the ATP10A/B/D group in mammals (11). Dnf2 is the primary plasma membrane GlcCer flippase in *S. cerevisiae*, and this protein transports the secondary and tertiary substrates PC and PE. Dnf1 also transports GlcCer, PC, and PE, but mostly localizes to intracellular compartments of the cell (10, 11, 20). A conserved Gln residue located in the middle of transmembrane segment 4 is critical for GlcCer transport, and Asn substitutions are sufficient to ablate GlcCer translocation in human and yeast enzymes without substantially altering recognition of glycerophospholipid substrates (11). Dnf1 and Dnf2 have been shown to be involved in sphingolipid homeostasis through their regulation by the flippase protein kinases (Fpk1/2) (21). Dnf1 and Dnf2 require phosphorylation by Fpk1/2 in the consensus motif R-X-S-ϕ-D/E for activity (22, 23). Fpk1/2 supports metabolic responses to changes in sphingolipid homeostasis through their connection to the TORC2 signaling network with yeast orthologs of PKD1 (Pkh1) and SGK1 (Ypk1). Most fungi produce both GlcCer and inositolphosphorylceramide (IPC), but *Saccharomyces cerevisiae* lacks GlcCer synthase and does not synthesize this lipid (24). IPC can be further modified with a second inositolphosphate group and/or mannose within the lumen of the Golgi to form MIPC and M(IP)_2_C, lipids that occupy the plasma membrane extracellular leaflet (25–27).

The plasma membrane has a tremendous diversity of lipid species, and P4-ATPases must sort through these lipids to establish membrane asymmetry. The identification and selection of which lipids to translocate and which to leave behind could be modulated by the composition of the membrane environment. The exofacial membrane leaflet consists of various lipid molecules which could be either high affinity or low-affinity substrates, potential inhibitors of transport, or allosteric modulators of the P4-ATPases. In our previous studies, we showed that unlabeled glucosylsphingosine (GlcSph) and lysoPC, lipids lacking a fatty acyl chain, could inhibit NBD-GlcCer transport by Dnf2 (11). These results suggested that the two substrate lipids are competing for a common transport pathway in the P4-ATPase or that changing the membrane composition influences flippase activity. In this study, we find that acute inhibition of sphingolipid synthesis or application of exogenous lipids modulates flippase substrate specificity.

## Results

### Acute inhibition of sphingolipid synthesis alters the substrate preference of Dnf1 and Dnf2

*S. cerevisiae* is part of a small clade of fungi that have evolutionarily lost the ability to make GlcCer and yet have retained GlcCer transporters (11, 24). Thus, Dnf1 and Dnf2 may facilitate uptake of GlcCer from exogenous sources (28–30). We hypothesized that inhibition of endogenous sphingolipid synthesis might enhance GlcCer uptake as a compensatory mechanism. Myriocin blocks sphingolipid biosynthesis at the first step in this pathway by inhibiting serine palmitoyl transferase (SPT) (31), while aureobasidin A (AbA) inhibits the inositol phosphorylceramide (IPC) synthase encoded by the *AUR1* gene (32) (Figure 1A). Wild type (WT), *dnf1*Δ, *dnf2*Δ, *fpk1,2*Δ (flippase protein kinase) strains were treated with myriocin and aureobasidin A for 30 mins and the uptake of NBD-PC and NBD-GlcCer was measured by flow cytometry (Figure 1). A *dnf1,2*Δ strain was also treated with the NBD-lipids and background (transport-independent) fluorescence was subtracted.

**Figure 1:**
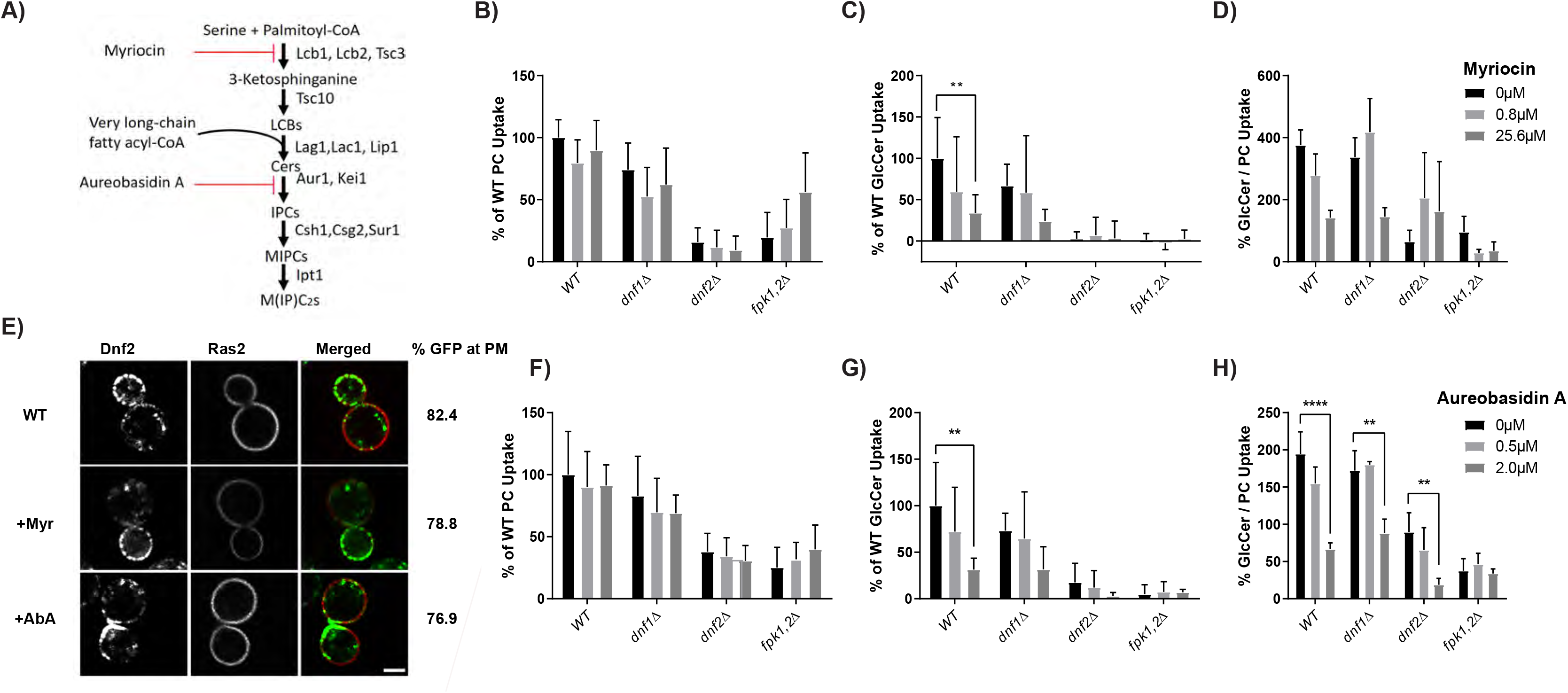
Acute inhibition of sphingolipid synthesis selectively reduces NBD-GlcCer transport. (A) Sphingolipid biosynthesis pathway in S. *cerevisiae* leading to the synthesis of IPC, MIPC and M(IP)_2_C including steps inhibited by myriocin and aureobasidin A. (B) Yeast strains BY4741 (WT), *dnf1*Δ and *dnf2*Δ strains were grown to the mid-log phase and treated with different concentrations of the myriocin for 30 mins at room temperature. Myriocin treated cells were incubated with NBD-PC or NBD-GlcCer for 30 mins and lipid uptake was measured using flow cytometry. Lipid uptake activities were plotted as the percentage of NBD-PC or NBD-GlcCer uptake for mock treated (0μM) wild-type cells. The GlcCer/PC ratio provides a measure of substrate preference. (C) Yeast strains BY4741, *dnf1*Δ, *dnf2*Δ and *fpk1*,2 strains were grown to the mid-log phase and treated with different concentration of aureobasidin A for 30 mins at room temperature. Aureobasidin A treated cells were incubated with NBD-PC or NBD-GlcCer. Lipid uptake was measured and plotted as in panel B. D) Wild type yeast strain expressing Dnf2-mNG and mCheery-Ras2 were treated with myriocin or aureobasidin A for 1 hr at room temperature. Cells were washed and resuspended in minimal media. Images were captured using the DeltaVision Elite Imaging System (GE Healthcare Life Sciences, Pittsburgh, PA) equipped with a 100 × objective lens followed by deconvolution using softWoRx software. Images were processed using ImageJ tool. Representative images are shown. (n=60) Scale Bar = 2um. Percentage of Dnf2-GFP at the plasma membrane are quantified by drawing circles inside and outside the plasma membrane using Image J to quantify the internal fluorescence and total fluorescence, respectively (n=60).

No significant change in uptake of NBD-PC was observed upon treatment with myriocin or aureobasidin A for any of the strains tested (Figure 1B, F). However, NBD-GlcCer uptake was selectively reduced in WT, *dnf1*Δ and *dnf2*Δ strains treated with myriocin and aureobasidin A (Figure 1C, G). Most of the plasma membrane transport activity is catalyzed by Dnf2 as indicated by the near ablation of transport in the *dnf2*Δ mutant. Interestingly, Dnf1 and Dnf2 retained a weak ability to transport NBD-PC in the *fpk1,2*Δ strain (~20% of WT transport activity in these experiments), but completely lost the ability to transport NBD-GlcCer. NBD-PC transport trended upward upon inhibition of sphingolipid synthesis in *fpk1,2*Δ but these differences were not significant. A measure of substrate preference for these flippases is described by the ratio of GlcCer transport to PC transport. The reduced GlcCer/PC uptake ratio in myriocin and AbA treated cells indicate that inhibition of sphingolipid biosynthesis specifically affects the GlcCer flippase activity of Dnf2 in a dose-dependent manner (Figure 1D, H). The localization of mNeonGreen (mNG) tagged Dnf2 to the plasma membrane, marked with Ras2-mCherry, was unaffected by inhibiting sphingolipid synthesis (Figure 1E). Thus, rather than enhancing GlcCer uptake by Dnf1 and Dnf2, inhibition of sphingolipid synthesis alters the substrate preference of these flippases by selectively reducing GlcCer transport.

### Sphingolipid biosynthesis mutants influence the substrate specificity of flippases

The major sphingolipids in *S. cerevisiae* (IPC, MIPC, M(IP)_2_C) are synthesized from ceramide by Aur1, Csg2, Csh1, Sur1 and Ipt1 (Figure 1A) (33–35). We tested whether knocking out nonessential biosynthetic genes required to produce mature glycosphingolipids would alter substrate specificity. All sphingolipid biosynthetic mutants exhibited a reduced uptake of NBD-PC and NBD-GlcCer (Figure 2A, B). In comparison with NBD-PC, these mutants displayed a greater reduction in flippase activity for NBD-GlcCer and therefore a reduced GlcCer/PC transport ratio (Figure 2C). The change in substrate preference could be partly explained by a greater degree of Dnf1 localization to the plasma membrane of the sphingolipid mutants, as this flippase transports NBD-PC and NBD-GlcCer at near equal rates (11). However, we found no change in localization of GFP-Dnf1 or Dnf2-mNG in sphingolipid biosynthesis mutants relative to WT cells (Figure 2D, E, Supplemental Figure S1). Thus, chronic loss of mature sphingolipids caused a reduction in Dnf1/2 activity and a change in substrate preference towards PC without altering localization of these proteins

**Figure 2:**
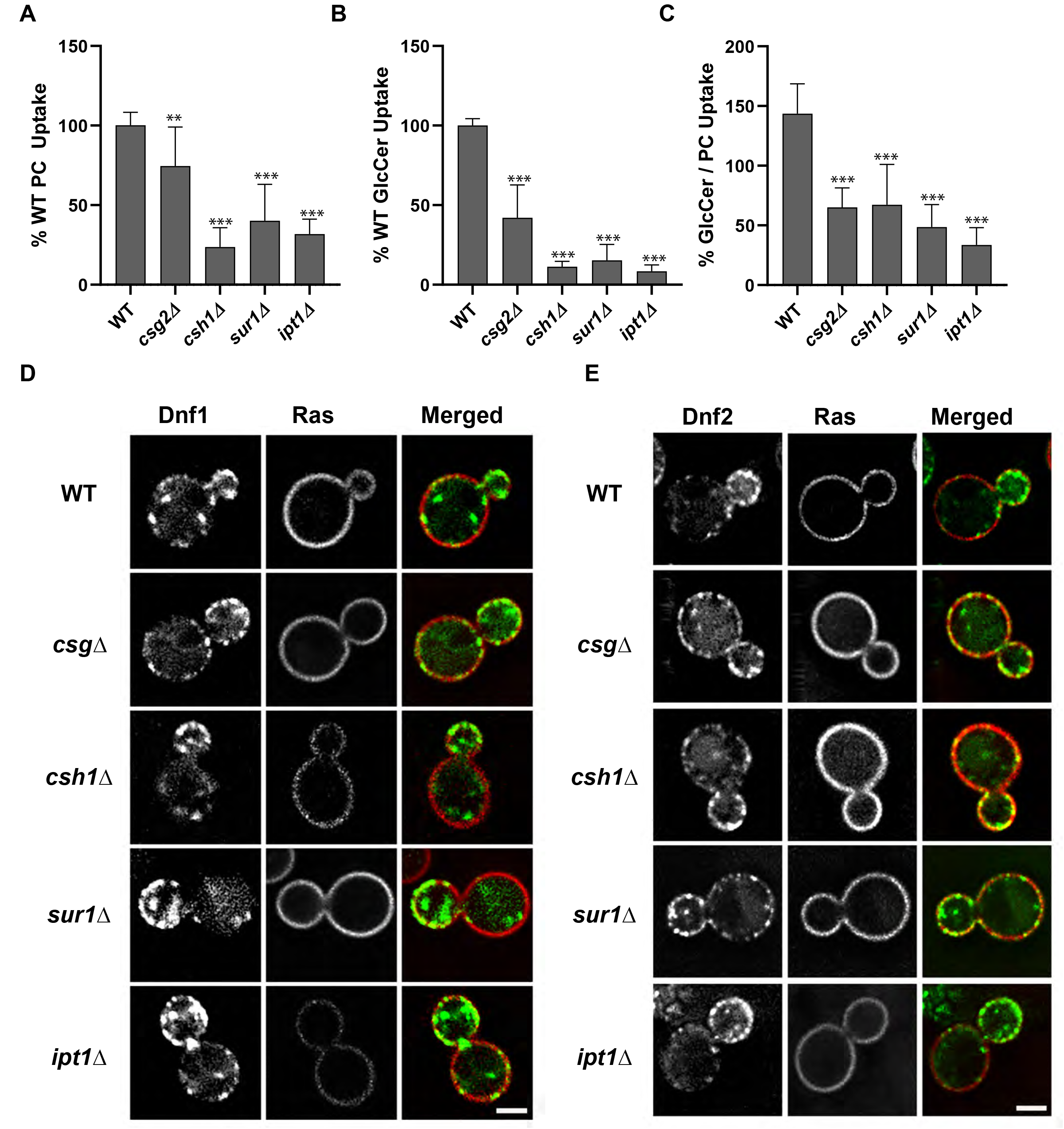
Perturbations to sphingolipid biosynthesis reduces plasma membrane flippase activity and alters substrate preference. Yeast strains BY4741 (WT), *dnf1*,2Δ *csg2*Δ, *csh1*Δ, *sur1*Δ, and *ipt1*Δ were grown to the mid-log phase and incubated with NBD-PC or NBD-GlcCer for 30 min. Lipid uptake activities were plotted as the percentage of NBD-PC (A) or NBD-GlcCer (B) uptake for wild-type cells, and the GlcCer/PC ratio (C). Statistical variance was tested using one-way ANOVAs test and Tukey’s post hoc analysis. * indicates p < 0.05, ** p< 0.01, *** p< 0.001. Error bars, S.D (n ≥ 9). (D) Strains BY4741 (WT), *csg2*Δ, *csh1*Δ, *sur1*Δ and *ipt1*Δ, expressing Dnf1 tagged with GFP and plasma membrane marker Ras2-mCherry were grown to mid-log phase in SD-Leu, URA media. Cells were washed and resuspended in minimal media for imaging. Images were captured using the DeltaVision Elite Imaging System (GE Healthcare Life Sciences, Pittsburgh, PA) equipped with a 100× objective lens followed by deconvolution using softWoRx software. Images were processed using ImageJ tool. Representative images are shown. Scale Bar = 2um (E) Yeast strains expressing Dnf2 tagged with mNG and Ras2-mCherry were grown to mid-log phase in SD-Leu media. Cells were washed and resuspended in minimal media for imaging. Images were processed using ImageJ tool. Representative images are shown. Scale Bar = 2um

Eisosomes are plasma membrane subdomains that play important roles in regulation of plasma membrane organization and have been linked to sphingolipid metabolism, the Pkh/Ypk signaling hub, and Slm1/Slm2 membrane stress sensors (Supplemental Figure S2). Therefore, we also tested eisosome structural component mutants (*pil1*Δ and *sur7*Δ) along with *slm1*Δ and *slm2*Δ to determine if these perturbations affect flippase activity or substrate specificity. We did not observe a significant difference in uptake of NBD-PC or NBD-GlcCer in *slm1*Δ or *pil1*Δ strains relative to WT, and *slm2*Δ showed a small decrease in both activities. Interestingly, uptake of NBD-PC was specifically reduced in the *sur7*Δ strain while NBD-GlcCer transport was unchanged (Figure 3A, B, C). We further examined eisosome structure in the sphingolipid synthesis mutants but found no difference in Sur7-RFP localization or appearance (Figure 3D). Therefore, these observations collectively suggest that eisosomes are not strong mediators of metabolic changes linked to flippase activity, but *sur7*Δ uniquely enhanced substrate preference towards GlcCer.

**Figure 3:**
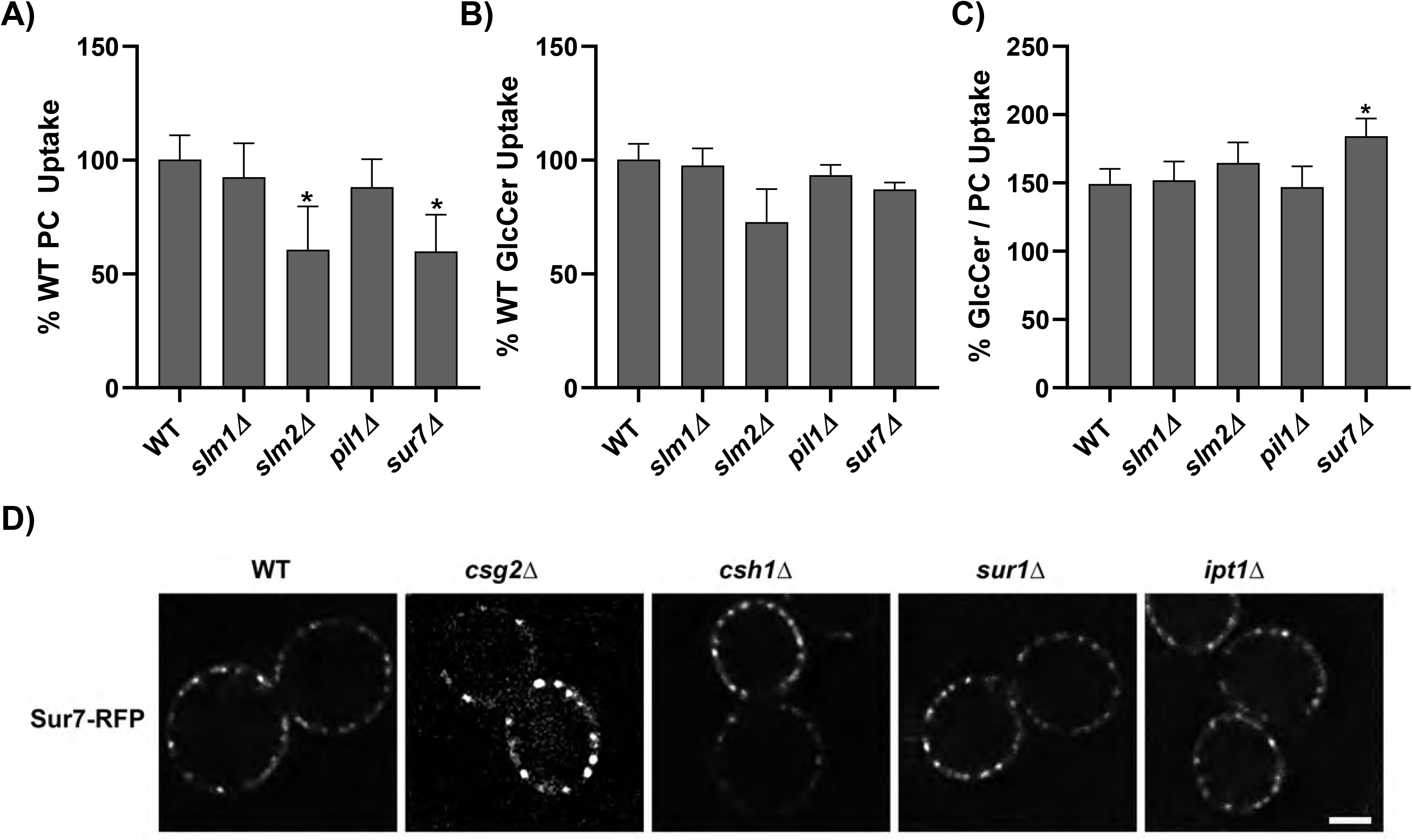
Influence of Eisosome mutants on flippase activity. Yeast strains BY4741(WT), *dnf1,2*Δ *slm1*Δ, *slm2*Δ, *pil1*Δ and *sur7*Δ Δ were grown to the mid-log phase and incubated with NBD-PC or NBD-GlcCer for 30 min. Lipid uptake activities were plotted as the percentage of NBD-PC (A) or NBD-GlcCer (B) uptake for wild-type cells, and the GlcCer/PC ratio (C). Statistical variance was tested using one-way ANOVAs test and Tukey’s post hoc analysis. * indicates p < 0.05. Error bars, S.D(n ≥ 9). (D)Yeast strains BY4741 (WT), *csg2*Δ, *csh1*Δ, *sur1*Δ, *ipt1*Δ, and *fpk1,2*Δ expressing Sur7 tagged with RFP were grown to mid-log phase in SD-Leu media. Cells were washed and resuspended in minimal media for imaging. Images were processed using ImageJ tool. Representative images are shown. Scale Bar = 2um

### Mutation of Fpk phosphorylation sites in Dnf2 to alanine ablates flippase activity

Inhibition of sphingolipid biosynthesis may influence flippase activity by reducing flippase protein kinase (Fpk1 and Fpk2) activity (Supplemental Figure S2). Dnf1 and Dnf2 are phosphorylated by Fpk1,2 and their phosphorylation strongly upregulates flippase activity (21), although the influence on GlcCer transport activity has not been assessed. We hypothesized that Fpk1,2 may be changing Dnf1 and Dnf2 substrate specificity through differential phosphorylation of these flippases (21, 22). Dnf2 possesses 5 Fpk1,2 phosphorylation sites of the consensus sequence R-X-S-L-D (Supplementary figure S3) (22). We mutated the serine residues in all five sites (5S-A, 5S-D) to prevent or mimic consensus Fpk1,2 phosphorylation. We sought to produce a gain-of-function *DNF2-5SD* allele to distinguish effects of phosphorylation versus more direct effects of membrane composition on Dnf2 activity. The 5S-A mutations resulted in loss of Dnf2 transport activity for both NBD-GlcCer and NBD-PC, without any change in flippase substrate specificity, implying that these residues are critical for flippase activity. Conversely, the Dnf2(5S-D) variant was functional and displayed equivalent transport activity to WT Dnf2 (Figure 4A). Yet despite the robust activity of the phospho-mimetic Dnf2, this allele could not suppress the lipid transport defects of the *fpk1,2* null strain (Figure 4B). Therefore, *DNF2(5S-D)* does not appear to be a gain-of-function allele capable of bypassing a flippase protein kinase deficiency and we did not further test this variant in the sphingolipid synthesis mutants.

**Figure 4:**
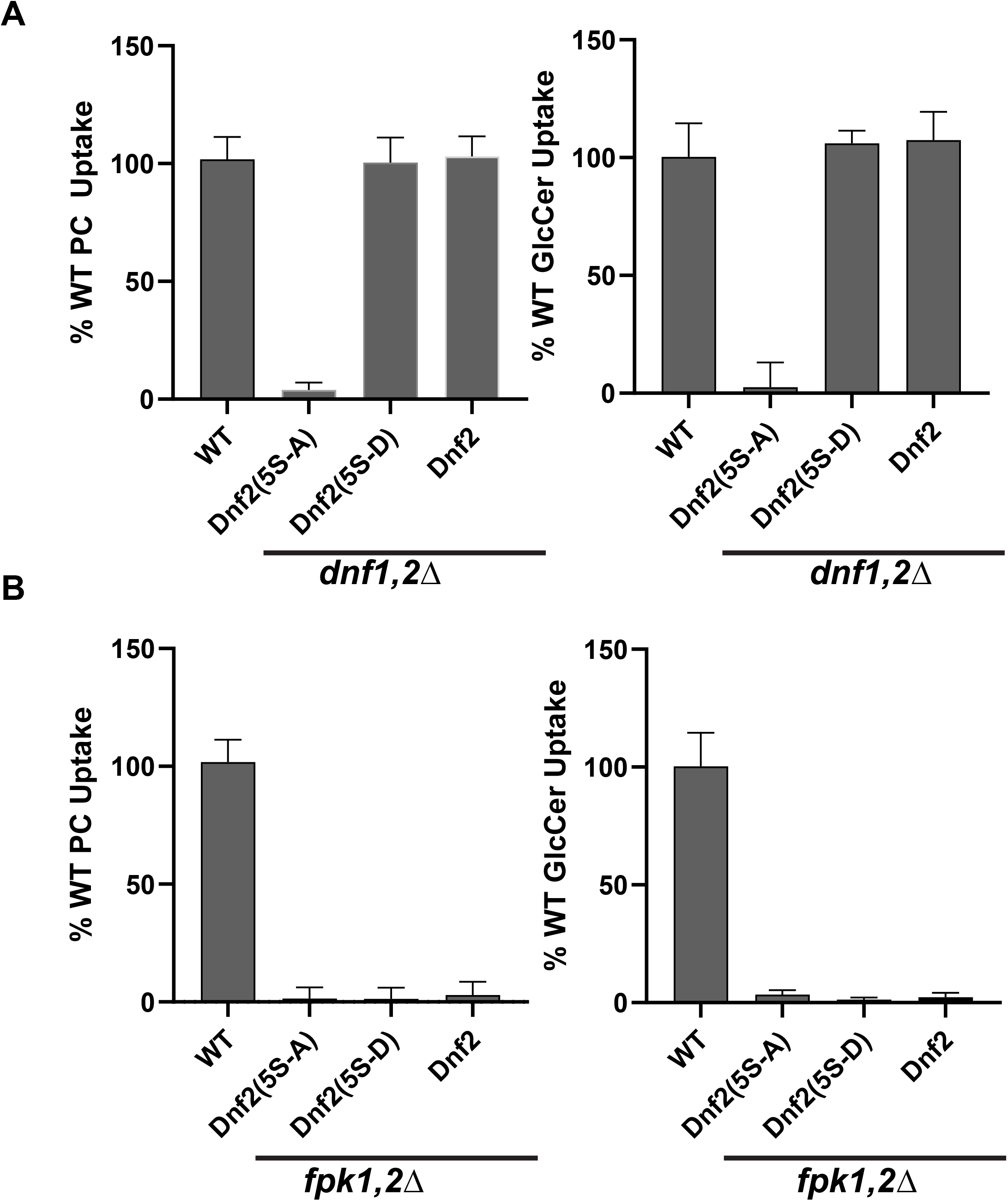
Fpk phosphorylation sites in Dnf2 are required for both NBD-PC and NBD-GlcCer transport activity. NBD-PC and NBD-GlcCer uptake was assayed in *dnf1,2*Δ (A) and *fpk1,2*Δ (B) strains harboring pRS313, pRS313-Dnf2(5S-5A), pRS313-Dnf2(5S-5D), and pRS313-Dnf2 plasmids. Lipid uptake activities were plotted as the percentage of NBD-PC and NBD-GlcCer uptake for wild-type cells and GlcCer/PC ratio. (n ≥ 9) ± S.D.

### Exogenous administration of unlabeled lipids alters Dnf2 lipid transport and substrate preference

The prior experiments indicated acute and chronic loss of endogenous sphingolipids altered the substrate specificity of Dnf2 by specifically decreasing GlcCer transport. These shifts in substrate preference were not driven by changes in intracellular signaling associated with the eisosome. We previously showed that unlabeled glucosylsphingosine (GlcSph, which is GlcCer without the N-acyl chain) and lysoPC could inhibit NBD-GlcCer transport by Dnf2 (11). Additionally, recent cryo-EM structures of the P4-ATPase family have identified only one lipid transport pathway through these enzymes (36–38). These results suggested these two lipid substrates are competing for a common transport pathway in the P4-ATPase.

Here, we sought to further address how acute application of exogenous lipids to the outer leaflet of the plasma membrane influences Dnf2 activity. NBD-labeled substrates were used to measure lipid transport, and unlabeled substrate (lyso-PC and GlcSph) and non-substrate lipids (sphingosine-1-phosphate – S1P, lactosylsphingosine – LacSph, lysosphingomyelin – lysoSM, and mono-acyl glycerol – 10MAG) (Figure 5A) were exogenously applied to determine their influence on substrate uptake and enzyme preference (Figure S4A). Lyso-lipids were chosen as potential inhibitors because Dnf2 appears to prefer these substrates (39). NBD-PC transport by Dnf2 was most sensitively inhibited by lyso-PC, GlcSph and LacSph (IC_50_ ~9-15μM). The inhibition by LacSph was surprising because NBD-LacCer is not a transport substrate (11). In contrast, lyso-SM and S1P inhibited NBD-PC transport weakly with an IC_50_ of approximately 40μM (Figure 5B).

**Figure 5.**
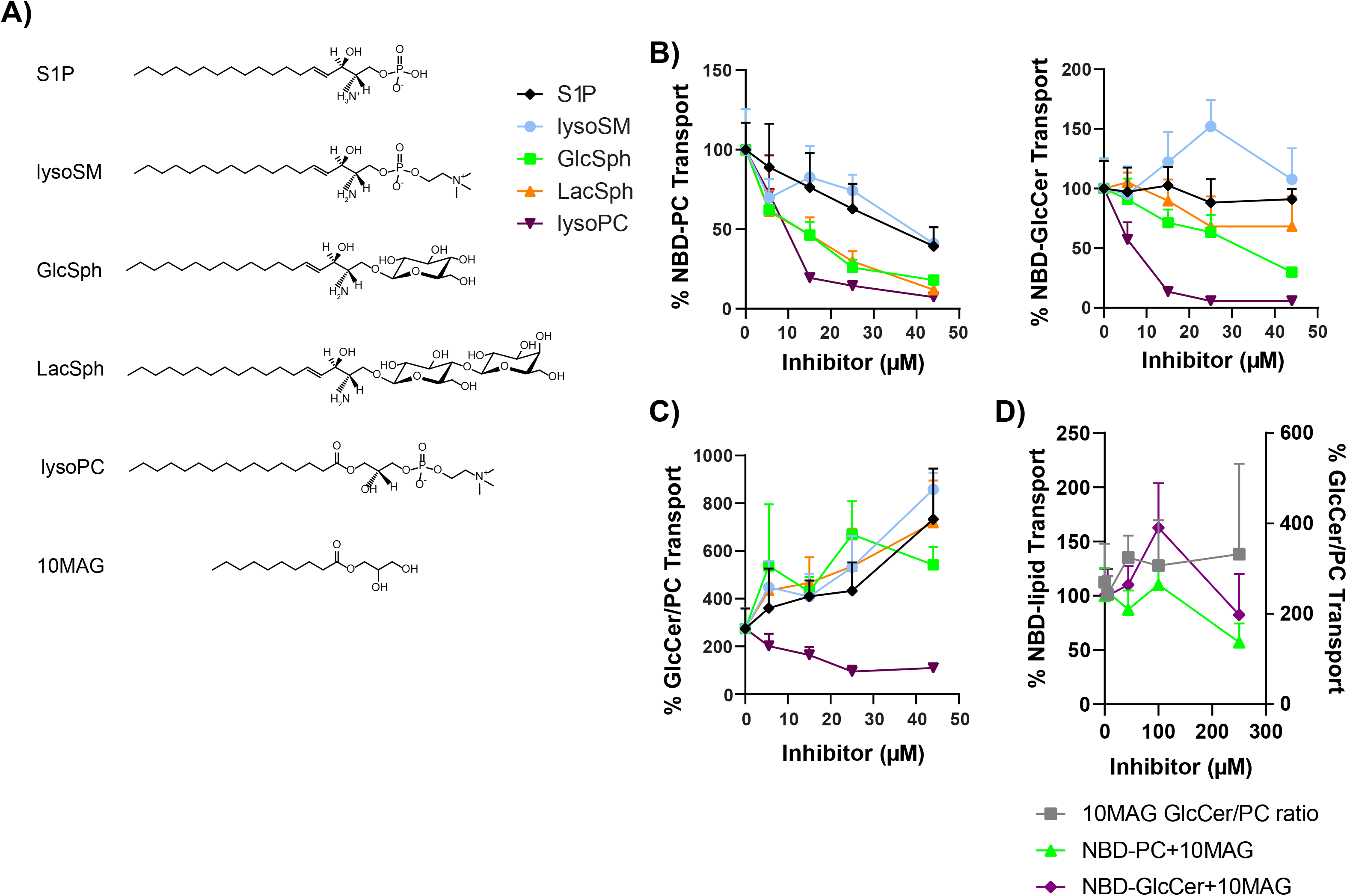
Exogenous administration of unlabeled lipids alters Dnf2 lipid transport and substrate preference. The chemical structures of all unlabeled lipids are presented (A). An increasing amount of phospholipids and sphingolipids were co-administered with NBD-PC or NBD-GlcCer to *dnf1*Δ *dnf2*Δ cells with *pRS313-Dnf2* or *pRS313*. Substrate transport was plotted relative to vehicle control (0μM) (B). The transport data were then transformed to reflect GlcCer/PC ratios over the same concentration ranges (C). A lipid detergent control, 10MAG, was tested for NBD-lipid transport inhibition up to 250uM (D), with NBD-lipid transport displayed on the left Y-axis, and the GlcCer/PC ratio displayed on the right Y-axis. Data presented is an XY scatter of the mean ± SD with connecting lines, (n≥6). Note: some of NBD-GlcCer inhibition data with lysoPC, GlcSph, and lysoSM used in (B) were previously used for Figure S2E in Roland et al., 2019.

Unexpectedly, the pattern of NBD-GlcCer transport inhibition by these lipids was different as compared to their influence on NBD-PC transport. Lyso-PC inhibited NBD-GlcCer transport efficiently (IC_50_ ~ 8μM) but a four-fold higher concentration of GlcSph was required for inhibition and LacSph weakly inhibited transport over the range of concentrations tested. Lyso-SM actually stimulated NBD-GlcCer transport at low concentrations and inhibited transport at high concentrations (Figure 5B, see (11) for the higher concentrations). These data were transformed as a ratio of NBD-GlcCer transport relative to NBD-PC to graphically show that these exogenous administrations of lipids altered the substrate preference of Dnf2, with most lipids inhibiting PC transport to a greater extent than GlcCer (Figure 5C).

To determine whether these preferential changes in lipid transport may be due to detergent-like effects of the exogenously administered lipids, we repeated these experiments using a mono-acyl glycerol with a C10 chain in the *sn*1 position (10MAG) (Figure 5D). Increasing co-administrations with 10MAG did not elicit substantial NBD-lipid transport inhibition up to 250uM (Figure 5D, left axis), and did not significantly change the preference of the enzyme (Figure 5D, right axis). Co-administration of unlabeled lipid with NBD-lipids may also sequester the probe in micelles away from the cells. We measured the influence of P4-ATPase expression, BSA back-extraction, and lipid co-administration on the accumulation of NBD-PC and NBD-GlcCer (Figure S4B-C). We found that lipid uptake in these cells is primarily driven by P4-ATPase activity, and additions of the unlabeled lipids did not significantly change the fluorescent signal of cells lacking Dnf1 and Dnf2, suggesting that the unlabeled lipids were not interfering or facilitating lipid administration or back-extraction (Figure S4B-C). These results support the conclusion that i) the exogenous administration of unlabeled lipids are capable of inhibiting NBD-lipid transport, ii) that the NBD-PC and NBD-GlcCer transport pathways can be differentially effected by its lipid environment, and iii) changing the lipid composition of the membrane can alter the observed preference of the lipid transporter.

Mammalian, plant, and fungal P4-ATPases are known to exhibit diverse substrate specificity, ranging from PS, PC, GlcCer, GalCer, PE, and SM (40). Several primary structural determinants have been mapped, suggesting how the enzyme may coordinate their amphipathic substrate(s) across the membrane (11, 36, 41–46). These studies have yielded numerous mutant P4-ATPase variants with gain-of-function or loss-of-function substrate specificities. We predicted that if the exogenous administrations of unlabeled lipids were having a direct impact on P4-ATPase substrate recognition or transport, the inhibitory properties of these lipids would track with the known substrate specificities of these mutant P4-ATPase variants.

A key structural feature of the P-type ATPase family is an entirely conserved Pro within the TM4 segment of the transmembrane domain. We have previously established that a conserved Gln residue located 4 residues N-terminal to this Pro (P – 4 Gln) is a conserved determinant of GlcCer transport by human and fungal P4-ATPases. The *DNF2^Q655N^* mutation dramatically reduces NBD-GlcCer transport, yet maintains substantial NBD-PC activity (11) (Figure S5), thereby indicating that this residue plays a critical role in glycosphingolipid recognition.

We expressed the *DNF2^Q655N^* variant in a *dnf1,2Δ* cell background and measured how four different unlabeled lipids altered its substrate transport profiles. We tested two classes of unlabeled lipids, substrate analogs and non-substrate analogs. We found that *DNF2^WT^* and *DNF2^Q655N^* enzymes responded similarly to lysoPC, lysoSM, GlcSph, and LacSph inhibition when translocating the NBD-PC substrate (Figure 6A). However, the *DNF2^Q655N^* mutation enhanced the potency of all unlabeled lipids to inhibit NBD-GlcCer transport (Figure 6B). Note that the data are normalized to the transport activity of each variant in the absence of the inhibitors. These data support the conclusion that the Q655N mutation specifically influences the ability of the Dnf2 enzyme to recognize and translocate the NBD-GlcCer substrate. Further, the enhanced potency of the unlabeled lipids for inhibiting NBD-GlcCer transport suggests that the enzyme is capable of recognizing exofacial lipid content, but that translocation of NBD-GlcCer through the enzyme may be impaired.

**Figure 6.**
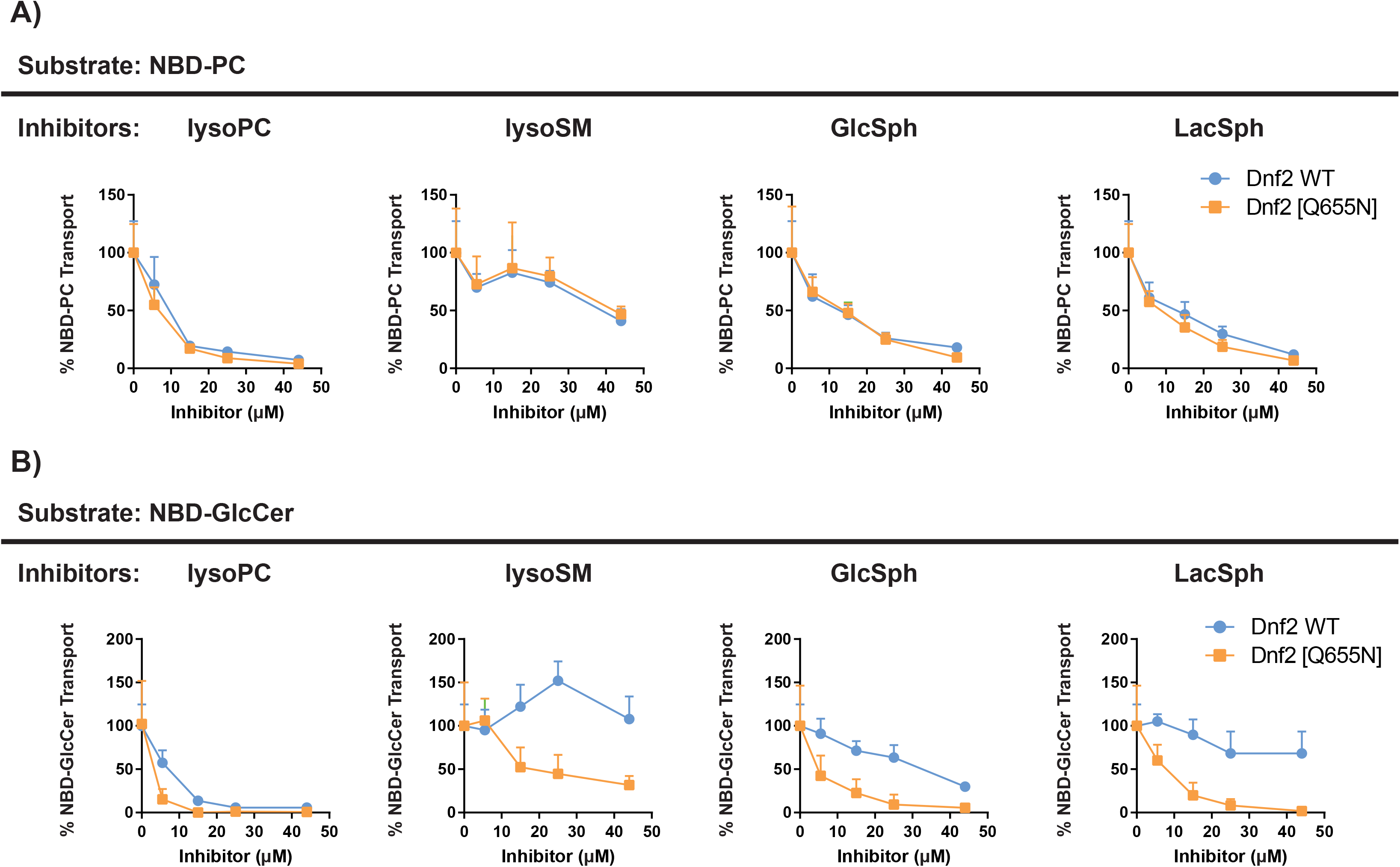
Dnf2^Q655N^ selectively alters NBD-GlcCer inhibition profiles. The *DNF2^WT^* and *DNF2^Q655N^* variants were tested in parallel for their ability to transport NBD-PC (A) and NBD-GlcCer (B) when co-administered with unlabeled lipids. NBD-lipid transport on the Y-axis is presented as a percentage of 0 μM lipid (vehicle). Data presented are mean ± S.D. (n≥6). Note: *DNF2 WT* NBD-GlcCer inhibition data with lysoPC, GlcSph, and lysoSM in (A) were previously published in Roland et al., 2019.

We have previously identified an N220S TM1 substitution in Dnf1 that gains the capacity to translocate NBD-SM. This mutation was isolated from a forward genetic screen for mutations that facilitated NBD-SM transport and reduced NBD-PC uptake (42). The homologous Asn in Dnf2 is residue 258, and Dnf2^N258S^ recapitulates this NBD-SM gain-of-function while still reducing NBD-PC uptake (Figure S5A, B, D). We tested whether Dnf2^N258S^ may impact NBD-PC transport or inhibition in the presence of increasing concentration of unlabeled lipids. Interestingly, NBD-PC inhibition via lysoSM was the only transport response that was significantly altered relative to WT (Table S1, Figure S6). The N258S mutation reduced the potency of lysoSM when administered with NBD-PC but not NBD-GlcCer, corroborating our previous studies of these mutants.

## Discussion

We report that the substrate preference of Dnf1 and Dnf2 is modulated by the composition of the membrane. Acute pharmacological inhibition of sphingolipid biosynthesis reduces GlcCer transport without altering PC transport, thus changing flippase substrate preference from glycosphingolipids to glycerophospholipids. Deletion of sphingolipid biosynthetic genes impacts transport of both substrates but again reduces GlcCer transport to a greater extent than PC transport. Conversely, application of exogenous sphingolipids to the outer leaflet, whether a substrate lipid or not, inhibits PC transport to a greater extent than GlcCer transport, whereas lysoPC inhibits both PC and GlcCer transport similarly. Finally, we find that mutations that alter substrate specificity of Dnf2 also alter its response to inhibitory lipids.

*S. cerevisiae* has retained flippases for GlcCer in spite of losing the ability to synthesize this lipid (24), perhaps to scavenge GlcCer from decaying fungal or plant material in the environment (28, 29). We hypothesized that the loss of endogenous sphingolipids would place a greater demand on the uptake of exogenous sphingolipids, and the cells would respond by upregulating Dnf1 and/or Dnf2 activity. This could provide an explanation for why the flippases are linked to a signaling network that also regulates sphingolipid synthesis (Figure S2). In addition, while IPC does not appear to be a transport substrate of Dnf1/2 (10) it seemed possible that this endogenous SL could be a competitive inhibitor of GlcCer transport, in which case blocking IPC synthesis should enhance GlcCer uptake. These possibilities, however, were not supported by our observations. As measured by NBD-PC uptake, no change in Dnf1/2 activity was observed upon acute inhibition of sphingolipid synthesis. Surprisingly though, NBD-GlcCer transport was reduced to 25 – 35% of that observed with mock-treated cells (Figure 1). Thus, it appears that the substrate preference was markedly changed such that these P4-ATPases now transported PC more efficiently than GlcCer. Chronic loss of mature sphingolipids induced by disruption of biosynthetic genes led to a small reduction in NBD-PC transport but a significantly larger impact on GlcCer transport and therefore a similar change in substrate specificity.

How would presence or absence of endogenous SLs influence the substrate specificity of these flippases? It is possible that modulation of substrate preference is an allosteric effect of the membrane environment on the P4-ATPase membrane domain where substrate is being flipped. Lipid is initially selected through the αβ subunit interface at the exofacial leaflet, it then docks within an entry gate site formed from M1 residues and the P – 4 Gln in M4, and transitions to an exit gate on the cytofacial surface where further selection of substrate can be elicited (11, 36, 41–43). Prior studies indicate conservative amino acid substitutions within this pathway can have a major influence on flippase substrate preference. In fact, several different mutations in M1, M4 and M6 cause similar changes to Dnf2 substrate specificity as a loss of sphingolipids (11, 41–43). The acute reduction of sphingolipids in the outer leaflet may alter the conformation of Dnf2 to specifically reduce affinity for GlcCer without perturbing its ability to bind PC. Similarly, the activities of the Ca^++^- and Na^+^/K^+^-ATPases is modulated by annular lipids that specifically associate with particular residues in this pump (47, 48). For Dnf2, non-substrate lipid induced conformational change could be a homeostatic mechanism where cells compensate for diminished IPC-derived glycosphingolipids by reducing GlcCer transport and leaving it in the outer leaflet. Whether this reflects a loss of specific interactions between the mature sphingolipids and Dnf2, or a general response to reduced lipid density in the outer leaflet is unclear.

Consistent with the idea that flippase substrate preference can be modulated in response to imbalances in lipid density between the two leaflets, we find that acute applications of exogenous sphingolipids, which would be expected to crowd the outer leaflet, preferentially inhibits NBD-PC transport much more than NBD-GlcCer transport (Figure 5). Thus, adding exogenous sphingolipid to the outer leaflet has the opposite effect on substrate preference as depleting endogenous sphingolipid. For NBD-PC transport, the nonsubstrate lipids S1P and lysoSM show a comparable dose-dependent inhibition with an IC_50_ value of approximately 45 μM while lysoPC and GlcSph substrates inhibit 3 – 4 fold more efficiently. It is possible that the SIP and lysoSM inhibition of NBD-PC transport is primarily due to crowding of the outer leaflet and greater potency is provided by substrates that can compete for the substrate binding sites. In this regard, LacSph is interesting because it is not a transport substrate yet inhibits NDB-PC transport comparably to GlcSph. This can be explained if LacSph can compete for entry gate binding but is too bulky to flip from entry to exit gate.

The pattern of NBD-GlcCer inhibition by exogenous lipids was unexpected. NDB-GlcCer is transported at twice the rate of NBD-PC in wild-type cells (11), implying GlcCer would have higher affinity for the flippases. However, lyso-PC is a significantly better inhibitor of NBD-GlcCer transport than is GlcSph. One possible explanation is that lysoPC has a comparable affinity for the entry gate but does not transition from entry to exit, or dissociate from the exit gate as efficiently as GlcCer. In this case, lysoPC would be better at tying up the enzyme and reducing enzyme turnover when assayed in competition for NBD-GlcCer transport. Another oddity was the stimulation of NBD-GlcCer transport by low concentrations of lyso-SM, where NBD-PC transport was weakly inhibited (Figure 5B). Similarly, this could be explained if lyso-SM, bearing the same phosphocholine headgroup as PC, could compete with endogenous lyso-PC for the entry gate but was not transported. This would reduce the number of enzymes bound unproductively to lyso-PC, perhaps holding it in the E2~P conformational state where NBD-GlcCer could efficiently displace lyso-SM for transport.

The involvement of the entry gate position in “sensing” membrane composition is supported by the observation that the Dnf2 [Q655N] mutation, which abrogates GlcCer transport much more dramatically than PC transport, also renders the remaining GlcCer transport much more sensitive to inhibition by other lipids (Figure 6). The strong reduction in GlcCer transport seen in the *DNF2^Q655N^* variant was also characterized by a strong increase in the potency of NBD-GlcCer inhibitors. These data demonstrated that the Q655N mutations selectively impacted the coordination of NBD-GlcCer, as the inhibitory profile of NBD-PC was unchanged relative to WT. Further, if binding or translocation of the NBD-GlcCer substrate were reduced, this could enhance the potency of any inhibitors. Thus, the observed reduction in NBD-GlcCer transport *in vivo* would be caused by the concerted reduction of GlcCer affinity and the increase inhibition from the enzymes’ lipid environment. Moreover, the Dnf2^N258S^ gain-of-function mutation was previously shown to increase the transport NBD-SM while reducing NBD-PC (11, 42). These effects were mirrored by changes in the potency of lysoSM inhibition of NBD-PC transport. However, the *reduced* potency of lysoSM as an NBD-PC inhibitor of the N258S mutant was unexpected; we had anticipated that the N258S mutant would be more susceptible to lysoSM inhibition if this were a competitive interaction. These data led us to consider that lysoSM may be allosterically modulating Dnf2 or altering lateral domain organization of the plasma membrane. Indeed, exogenous lysoSM administrations did increase WT transport of NBD-GlcCer to nearly 150% that of the vehicle treatment (Figure 6B). We also found that a short chain monoacylglycerol exerted no significant change in substrate specificity. These observations support the idea that we are observing specific changes in substrate preference in these studies as opposed to a more general response to membrane perturbation.

It is also possible that differential phosphorylation of Dnf1 and Dnf2 by the flippase protein kinases can modulate substrate specificity. Membrane stress induces changes in the localization of Slm1/2 proteins leading to activation of TORC2-Ypk1 signaling (49). Ypk1 directly phosphorylates Fpk1 and Fpk2 to inhibit their activity and conversely, Fpk1/2 phosphorylates Ypk1 to inhibit its phosphorylation activity (Figure S2) (22). Cellular status regulates the balance between Ypk and Fpk activity, and therefore flippase activity. Inhibition of sphingolipid biosynthesis stimulates TORC2 and thereby upregulates Ypk1 activity (49), which should inhibit Fpk1 and Fpk2 and cause a reduction of flippase activity (22). Indeed, we observe reduced PC and GlcCer flippase activity in mutants deficient for mature glycosphingolipids that could be caused by reduced phosphorylation of the flippases. We also see some indication that the *fpk1,2*Δ mutant displays a greater loss of GlcCer transport activity than PC transport (Figure 1). However, in other experiments we observed a complete loss of both PC and GlcCer activity in the *fpk1,2*Δ strain (Figure 4). The reason for these differences are unclear and may reflect subtle differences in nutrient status of the cells at the time they are assayed. We attempted to test the influence of phosphorylation by generating phosphomimetic mutations in Dnf2 (Dnf2(5S-D)) hypothesizing that this variant would be constitutively active and unresponsive to perturbation of sphingolipid synthesis. While a phosphorylation deficient Dnf2 mutant (5S-A) is completely devoid of flippase activity, the potential phosphomimetic Dnf2(5S-D) variant displays WT activity for both GlcCer and PC transport. However, Dnf2(5S-D) fails to display any activity in the *fpk1,2*Δ mutant. None of the single site phosphodeficient or phosphomimetic mutants showed changes in flippase activity (unpublished observation). These results might suggest that direct phosphorylation of Dnf2 is required for activity, but Fpk-dependent phosphorylation of other proteins are also a critical aspect of regulating flippase activity. However, the precise influence of these Ser to Ala and Ser to Asp mutations on Dnf2 structure/function are unknown and more work is needed to understand the complex regulation of flippase activity by protein kinases.

We hypothesized that specialized membrane compartments called eisosomes may play roles in sensing changes in membrane lipid composition that would be transduced to the flippases. Inhibition of sphingolipid biosynthesis can lead to accumulation of long-chain sphingoid bases (LCBs) capable of stimulating the eisosome-localized Pkh1,2 proteins (50). However, deletion of a major eisosome component, Pil1, does not alter flippase transport activity or substrate specificity. Deletion of the membrane sensor Slm2 does cause a small reduction in flippase activity but no change in specificity. Interestingly, the *sur7*Δ mutant shows an increase in the GlcCer/PC ratio caused by a preferential reduction in NBD-PC transport (Figure 3). Sur7 is a component of eisosomes and loss of this protein perturbs hydroxylation of sphingolipids by an unknown mechanism (51). The observed change in flippase substrate specificity in *sur7*Δ is similar to the effect of adding exogenous sphingolipid; perhaps *sur7*Δ causes crowding of the outer leaflet to give the similar change in flippase substrate preference.

The flippases are a unique set of transporters in that they actively modify the membrane itself rather than simply moving substrate across the membrane from one aqueous compartment to another. As such, flippases exist in a membrane environment of mixed substrate and putative inhibitors and/or allosteric modulators (Figure 7). Each lipid molecule has the potential to consistently sample the active site of the flippase, and their presence or absence could inherently impact the transport activity of the enzyme. We tested this hypothesis by changing the exofacial membrane composition through depletion of endogenous sphingolipids and exogenous lipid additions. We found that both substrates and non-substrates could influence the transport activity and preference of the flippases. Alterations to substrate transport and preference appears to be caused by the concerted influence of flippase phosphorylation, allosteric regulation, and competitive inhibition.

**Figure 7:**
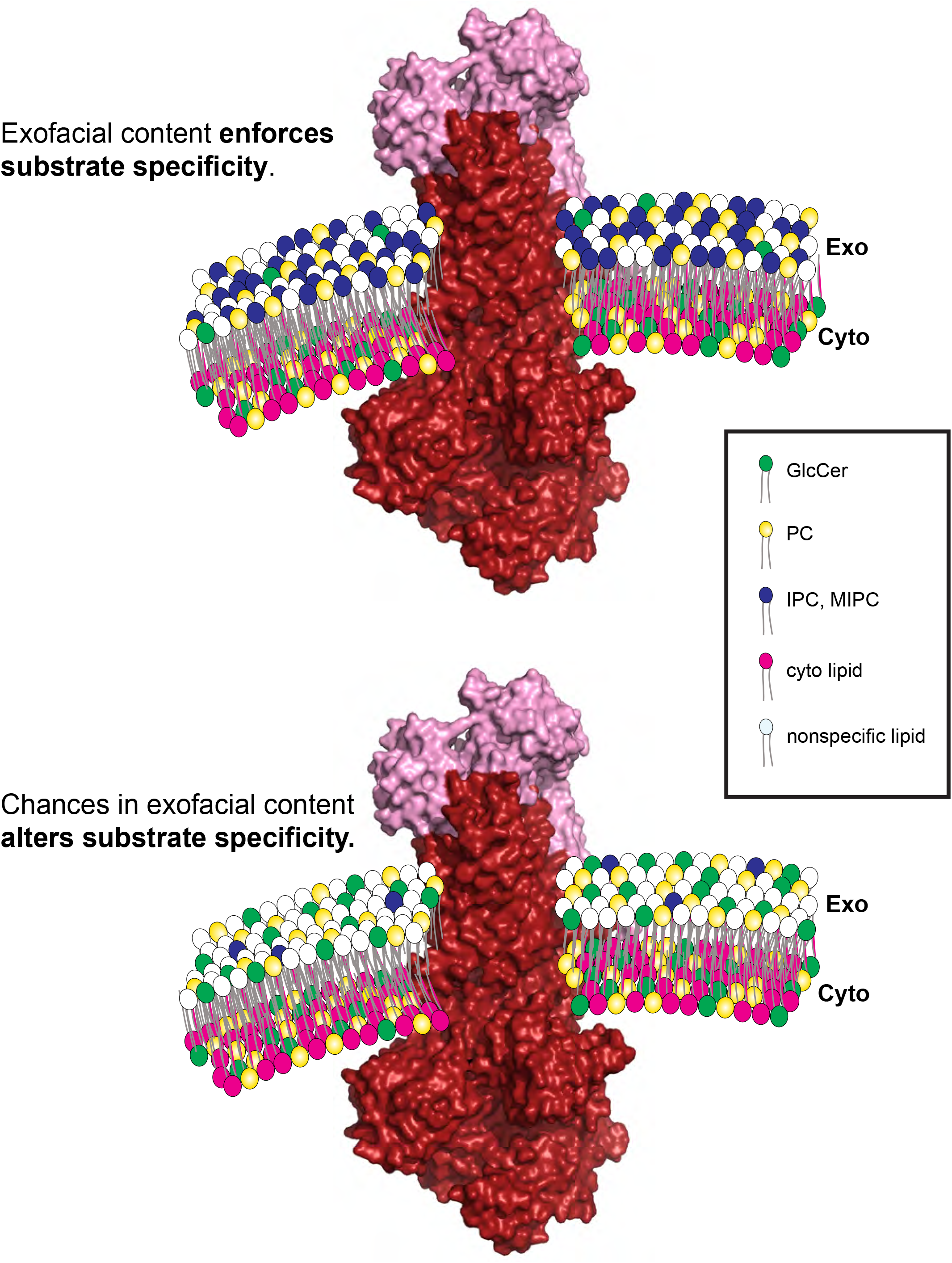
Exofacial lipid composition influences the flippase activity. The exofacial membrane leaflets consist of various lipid molecules which could be either high affinity or low-affinity substrates, potential inhibitors of transport, or allosteric modulators of the P4-ATPases. The exofacial composition enforces the substrate specificity or concedes secondary substrate transport. Figure was created using a model of the Drs2:CDC50 complex (PDB: 6ROI)(37).

## Materials and Methods

### Reagents

All yeast culture reagents were purchased from Sigma-Aldrich and BD scientific. All the lipids used in this study were purchased from Avanti Polar Lipids. Myriocin was from Cayman Chemicals while aureobasidin A was from Takara Bio.

### Strains and plasmid construction

Yeast strains and plasmid used in the study are listed in Table S2 and S3. Yeast strains were grown in YPD, SD, and selective media. Yeast strains were transformed using standard transformation techniques. Dnf2 was epitope-tagged on the C-terminus with mNeonGreen by a PCR-based genomic integration method using pFA6a-mNG-Hygro as the template (52). The plasma membrane marker mCherry-Ras2 was subcloned from mCherry-Ras2-YIplac204 (53) to pRS315 (54). To create DNA constructs, PCR amplified products were generated and two-or three-piece Gibson assembly was performed per manufacture’s instruction (New England Biolabs)

### Lipid Uptake assays

Lipid uptake assays were conducted as described previously (11, 42). Cells were grown to the mid-log phase and approximately 1 *A*_600_ of cells were collected for each strain. NBD-PC and NBD-GlcCer were dried and resuspended in 100% ethanol, then added to the designated ice-cold growth medium at a final concentration of 2 μg/ml. Final ethanol volumes were ≤0.5%. Cells were incubated on ice with the pre-chilled NBD-lipid+media for 30 min. Following incubation, the cells were washed twice with pre-chilled SA medium (SD medium + 2% (w/v) sorbitol + 20 mM NaN_3_) supplemented with 4% (w/v) fatty acid-free BSA. Finally, cells were resuspended in SA with 5 μM propidium iodide (PI) and NBD-lipid uptake was immediately measured by flow cytometry. The order in which samples were prepared and processed was varied in each experiment to reduce potential experimental error. Each experiment consisted of three independent biological replicates for each strain, repeated three times.

### Inhibition assays

Inhibitor experiments were performed as defined previously (11). An illustrated depiction of the method can also be found in Figure S1A. Briefly, the experiment was conducted as outlined above except the NBD-lipid media was not pre-chilled before cellular administration. The lipids + media were allowed to remain at room temperature to improve the solubility of the inhibitors during cell application. Lipid inhibitors were solubilized in 100% ethanol and added to SD supplemented with NBD-lipid probes at the indicated final concentrations. Solubility was closely monitored, and all inhibitor lipids solubilized easily in ethanol except LacSph, which required sonication and gentle heating at 37°C. Care was taken to ensure the final concentration of ethanol in all assay mixtures was 2.2%. No lipid insolubility was noted when lipid mixtures were added to media. Each experiment consisted of three independent biological replicates for each strain, repeated twice per inhibitor concentration.

### Myriocin and Aureobasidin A treatment

Myriocin and aureobasidin A experiments were conducted as outlined above, except the cells were pre-incubated with SD + the designated concentration of each compound at room temperature for 30 min. PI staining was not enhanced with this acute treatment (Figure S7). Each experiment consisted of three independent biological replicates for each strain, repeated three times.

#### Flow cytometry

All flow cytometry experiments were conducted on a three-laser BD LSRII (BD Biosciences) operating a FACS Diva 6.1.3 software package. Single-cell populations were identified by forward- and side-scatter as illustrated in Figure S7. PI was used to exclude dead cell populations and those with lost membrane integrity, and NBD signal was measured with FITC filters (530/30 nm band-pass filter with 525 nm long-pass filter). At least 10,000 cells were analyzed per experimental replicate.

#### Fluorescence microscopy

To visualize GFP or mCherry-tagged proteins, cells were grown to mid-logarithmic phase in YPD or minimal media. Cells were washed with fresh media for 3 times and resuspended in fresh SD media. Cells were then mounted on glass slides and observed immediately at room temperature. Images were acquired using a DeltaVision Elite Imaging System (GE Healthcare Life Sciences, Pittsburgh, PA) equipped with a 100 × objective lens followed by deconvolution using softWoRx software (GE Healthcare Life Science).

#### Image analysis and quantification

Image analysis was performed as previously described(55). Overlay images were created using the merge channels function of ImageJ software (National Institutes of Health). GFP-Dnf1 and Dnf2-GFP at the plasma membrane are quantified by drawing circles inside and outside the plasma membrane using Image J to quantify the internal fluorescence and total fluorescence, respectively. The internal fluorescence was subtracted from the total to give the GFP intensity at the plasma membrane. At least 60 randomly chosen cells from three biological replicates (independently isolated strains with the same genotype) were used to calculate the mean and standard deviation

#### Statistical analysis

The statistical analyses were performed using GraphPad Prism 8.4.1. Variance was calculated using one-way ANOVAs test and comparisons with wild type strains were tested with Tukey’s post hoc analysis. The P value represents the significance of the data * indicates p < 0.05, ** p< 0.01, *** p< 0.001.

## Data availability

The primary data will be available upon request. No material transfer agreements are required for data accessibility.

## Acknowledgments

We thank Graham lab members for helpful discussion during studies and Jordan T. Best for creating the *fpk1,2Δ* strain. We thank Kathy Gould for providing plasmids used in the studies. The yeast flow cytometry experiments were performed in the Vanderbilt Medical Center (VMC) Flow Cytometry Shared Resource.

## Funding and additional information

This work was supported by National Institutes of Health Grants R01-GM107978 (to T. R. G.) and F32-GM116310 (to B.P.R.). The VUMC Flow Cytometry Shared Resource is supported by NIH grants to the Vanderbilt Ingram Cancer Center (P30-CA68485) and the Vanderbilt Digestive Disease Research Center (P30-DK058404). The content is solely the responsibility of the authors and does not necessarily represent the official views of the National Institutes of Health.

